# Chasing perfection: validation and polishing strategies for telomere-to-telomere genome assemblies

**DOI:** 10.1101/2021.07.02.450803

**Authors:** Ann M. Mc Cartney, Kishwar Shafin, Michael Alonge, Andrey V. Bzikadze, Giulio Formenti, Arkarachai Fungtammasan, Kerstin Howe, Chirag Jain, Sergey Koren, Glennis A. Logsdon, Karen H. Miga, Alla Mikheenko, Benedict Paten, Alaina Shumate, Daniela C. Soto, Ivan Sović, Jonathan MD Wood, Justin M. Zook, Adam M. Phillippy, Arang Rhie

## Abstract

Advances in long-read sequencing technologies and genome assembly methods have enabled the recent completion of the first Telomere-to-Telomere (T2T) human genome assembly, which resolves complex segmental duplications and large tandem repeats, including centromeric satellite arrays in a complete hydatidiform mole (CHM13). Though derived from highly accurate sequencing, evaluation revealed that the initial T2T draft assembly had evidence of small errors and structural misassemblies. To correct these errors, we designed a novel repeat-aware polishing strategy that made accurate assembly corrections in large repeats without overcorrection, ultimately fixing 51% of the existing errors and improving the assembly QV to 73.9. By comparing our results to standard automated polishing tools, we outline common polishing errors and offer practical suggestions for genome projects with limited resources. We also show how sequencing biases in both PacBio HiFi and Oxford Nanopore Technologies reads cause signature assembly errors that can be corrected with a diverse panel of sequencing technologies

## INTRODUCTION

Genome assembly is a foundational practice of quantitative biological research with increasing utility. By representing the genomic sequence of a sample of interest, genome assemblies enable researchers to annotate important features, quantify functional data, and discover/genotype genetic variants in a population^1–6^. Modern draft eukaryotic genome assembly graphs are typically built from a subset of four Whole Genome Shotgun (WGS) sequencing data types: Illumina short reads^7,8^, Oxford Nanopore Technologies (ONT) long reads^9^, PacBio Continuous Long Reads (CLR), and PacBio High-Fidelity (HiFi) long reads^9,10^, all of which have been extensively described^7–10^. However, we note that even the high-accuracy technologies produce sequencing data with some noise caused by platform-specific technical biases that require careful validation and polishing^11,12,1,10,13^.

Current genome assembly software attempts to reconstruct an individual or mosaic haplotype sequence from a subset of the above WGS data types. Some assemblers do not attempt to correct sequencing errors^14^, while others attempt to remove errors at various stages of the assembly process^15–19^. Regardless, technology-specific sequencing errors usually lead to distinct assembly errors^13,20^. Additionally, suboptimal assembly of specific loci often causes small and large errors in draft assemblies^21,22^. The process of removing these errors from draft genome assemblies is known as “polishing”. Most polishing tools use an approach that is similar to sequence-based genetic variant discovery. Specifically, reads from the same individual are aligned to a draft assembly, and putative “variant”-like sequence edits are identified^22,23^. In diploids, heterozygous “alternate” alleles are interpreted as genuine heterozygous variants, while homozygous alternate alleles are interpreted as assembly errors to be corrected. Some polishing tools, such as Quiver/Arrow, Nanopolish, Medaka, DeepVariant, and PEPPER leverage specialized models and prior knowledge to correct errors caused by technology-specific bias^24–28^. Others, such as Racon^29^, use generic methods to correct assembly errors with a subset of sequencing technologies^29–31^. These generic tools can utilize multiple data types to synergistically overcome technology-specific assembly errors.

In the summer of 2020, the Telomere-to-Telomere (T2T) consortium convened an international workshop to assemble the first-ever complete sequence of a human genome. As heterozygosity can complicate assembly algorithms, the consortium chose to assemble the highly homozygous genome of a complete hydatidiform mole cell line (CHM13hTERT; abbr. CHM13). Primarily using HiFi reads and supplemented with ONT reads, the consortium built a highly accurate and complete draft assembly (CHM13v0.9) that resolved all repeats with the exception of the rDNAs and selected a locally haplotype-consistent path for the CHM13 genome^1^. CHM13v0.9 contained about 1 error in every 10.5 Mb (Q70.22), and while this was highly accurate by traditional standards, we, as part of the consortium, sought to correct all lingering errors and omissions in this first truly complete assembly of a human genome.

Here, we describe techniques developed to carefully evaluate the accuracy and completeness of a complete human genome assembly using multiple complementary WGS data types. Our evaluation of the initial draft CHM13 assembly discovered a number of assembly errors, and so we created a custom polishing pipeline that was robust to genomic repeats and technology-specific biases. By applying this polishing pipeline to CHM13v0.9, we made 1,457 corrections, replacing a total of 12,234,603 bp of sequence with 10,152,653 bp of sequence, ultimately leading to the landmark CHM13v1.1 assembly representing the first complete human genome ever assembled. Our edits increased the estimated quality value to Q73.94 while mitigating haplotype switches in the already-haplotype-consistent consensus sequence. Further, we extended the truncated p-arm of chromosome 18 to encompass the complete telomere, and polished all telomeres with a new specialized PEPPER-DeepVariant model. Our careful evaluation of CHM13v1.1 confirmed that polishing did not overcorrect repeats (including rDNAs) nor did it cause false-positive edits to protein-coding transcripts. Additionally, we identified a comprehensive list of putatively heterozygous loci in the CHM13 cell line, as well as sporadic loci where read alignments still indicated exceptionally low coverage. Finally, we uncovered common mistakes made by standard automated polishing pipelines and provide best practices for other genome assembly projects.

## RESULTS

### Initial evaluation of CHM13v0.9

The T2T Consortium has collected a comprehensive and diverse set of publicly available WGS sequencing and genomic map data (Illumina PCR-free, PacBio HiFi, PacBio CLR, and ONT) for the nearly-complete homozygous CHM13 cell line (https://github.com/marbl/CHM13). As part of the consortium, we drew upon these sequencing data to generate a custom pipeline **(Fig. 1)** to evaluate, identify and correct lingering errors in CHM13v0.9.

**Figure 1.**
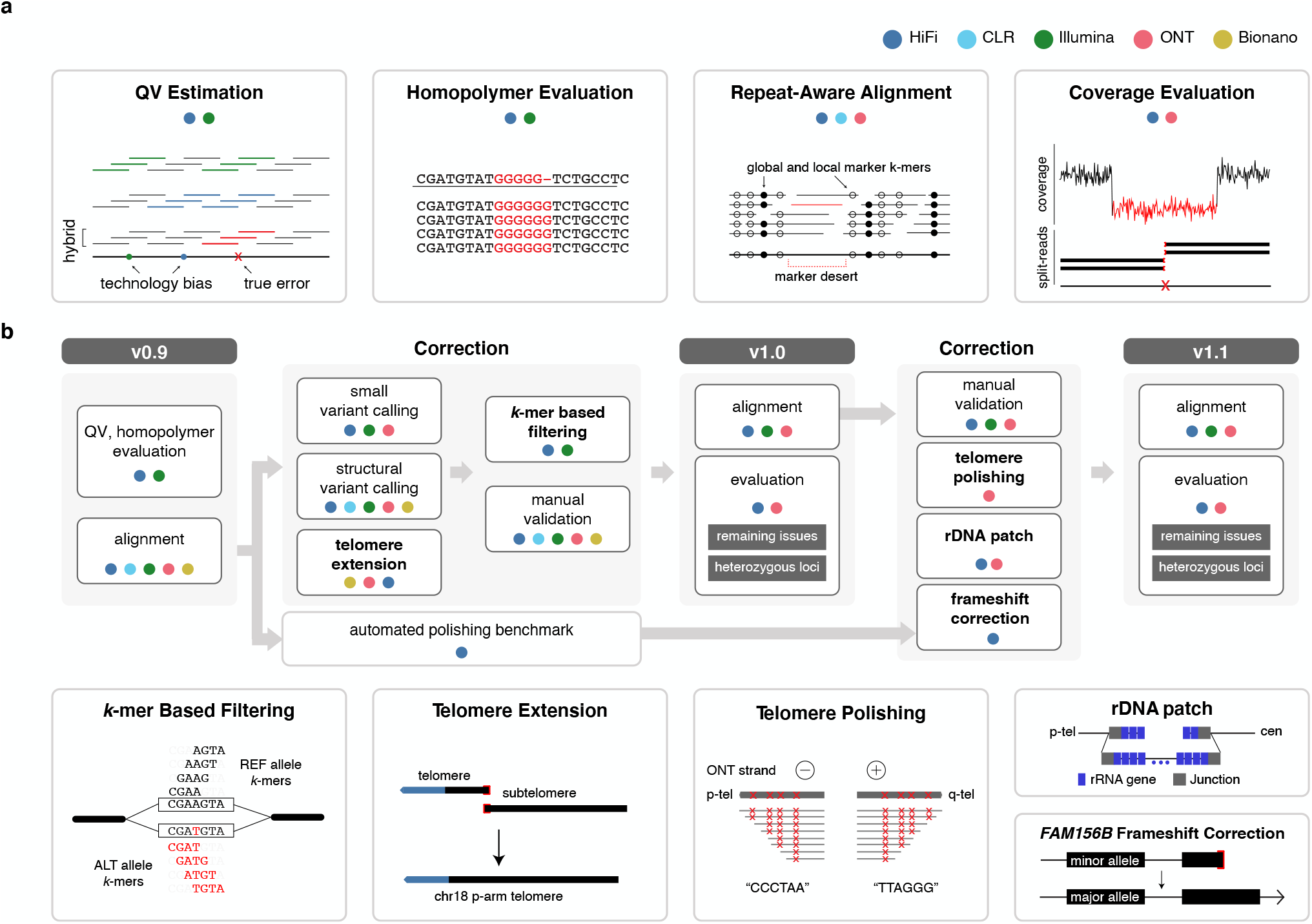
An overview of the evaluation and polishing strategy developed to achieve a complete human genome assembly. **a**, The evaluation strategies used to assess genome assembly accuracy before (CHM13v0.9) and after (CHM13v1.0 and CHM13v1.1) polishing. **b**, The “do no harm” polishing strategy developed and implemented to generate CHM13v1.0 and CHM13v1.1.

We first derived *k*-mer-based quality estimations (*k* = 21bp) of CHM13v0.9 using Merqury^32^ using both Illumina and HiFi reads. While estimating the Illumina reads QV, we found 15,723 *k*-mers present in the assembly and not the reads (erroneous *k*-mers), leading to an estimated base quality of Q66.09. Using HiFi reads, we found 6,881 error *k*-mers (Q69.68) (**Fig. 2a**). To test how technical sequencing bias may have influenced this QV estimation, we examined the *k*-mer multiplicity and sequence content of assembly *k*-mers absent from one technology but present in the other. Here, our results indicated that *k*-mers missing from Illumina reads were present with expected frequency in HiFi and were enriched for G/C bases. Conversely, *k*-mers missing in HiFi were present with higher frequency in Illumina reads with A/T base enrichment (**Fig. 2b**). However, we identified no particular enrichment pattern in the number of GA or CTs within the *k*-mers, possibly due to the short *k*-mer size chosen (**Supplementary Fig. 1a**). Most of the *k*-mers absent from HiFi reads were located in patches derived from a previous ONT-based assembly (CHM13v0.7), which were included to overcome regions of HiFi coverage dropout^1^(**Supplementary Fig. 1b-c**). These findings highlighted that platform-specific sequencing biases were underestimating the QV when measured from a single sequencing platform. To overcome this, we created a hybrid *k*-mer database that combined these platforms to be used for QV estimation (**Supplementary Fig. 1d**). Unlike the default QV estimation in Merqury, we removed low frequency *k*-mers to avoid overestimated QVs caused by excessive noise accumulated from both platforms. We estimated base level accuracy as Q70.22 with 6,073 missing *k*-mers **(Supplementary Table 1)**. We note that this estimate does not account for the rarer case of *k*-mers present in the reads but misplaced or falsely duplicated in the assembly.

**Figure 2.**
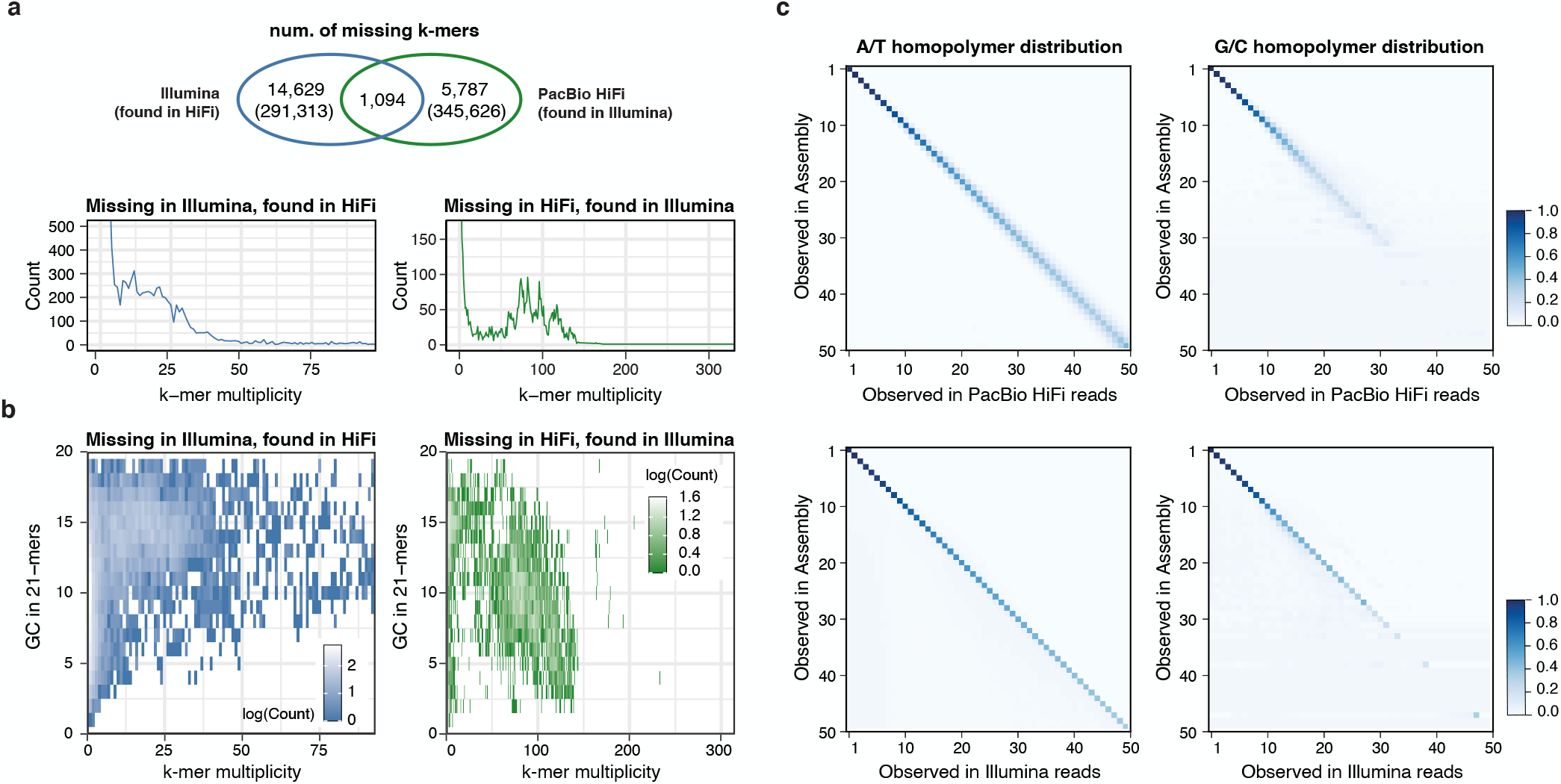
Sequencing biases in PacBio HiFi and Illumina reads. **a**, Venn Diagram of the “missing” *k*-mers found in the assembly but not in the HiFi reads (green) or Illumina reads (blue). Except for the 1,094 *k*-mers that were absent from both HiFi and Illumina reads, error *k*-mers were found in the other sequencing platform with expected frequency, matching the average sequencing coverage (lower panels). **b**, Missing *k*-mers from **a** with its GC contents, colored by the frequency observed. Low frequency erroneous *k*-mers did not have a clear GC bias. *k*-mers found only in HiFi had a higher GC percentage, while higher frequency *k*-mers tend to have more AT rich sequences in Illumina. **c**, Homopolymer length distribution observed in the assembly and in HiFi reads (upper) or Illumina reads (lower) aligned to that position. Longer homopolymers in the consensus are associated with length variability in HiFi reads especially in the GC homopolymers. The majority of the Illumina reads were concordant with the consensus.

Despite the high accuracy of CHM13v0.9 (Q70.22), we expected to find consensus sequence errors related to the systematic presence of homopolymer- or repeat-specific issues in HiFi reads^9,33^. To detect these, we generated self-alignments by aligning CHM13 reads to CHM13v0.9 for each WGS sequencing technology. Though each data type required technology-specific alignment methods (**Methods**), we highlight our use of Winnowmap2 that enabled robust alignment of long-reads to both repetitive and non-repetitive regions of CHM13v0.9^34,35^. To understand the homopolymer length differences between the assembly and the reads, we derived a confusion matrix from Illumina read alignments showing discordant representation of long homopolymers between the Illumina reads and the assembly (**Fig. 2c)**. Altogether, the QV and homopolymer analysis suggested that CHM13v0.9 required polishing to maximize accuracy of a complete human genome.

### Identification and correction of assembly errors

To address assembly flaws identified during evaluation, we aimed to establish a customized polishing pipeline that would avoid false positive polishing edits and maintain local haplotype consistency (**Fig. 1b**). We identified and corrected small errors (<=50bp) using several small variant calling tools from self-alignments of Illumina, HiFi and ONT reads to CHM13v0.9. To call both single-nucleotide polymorphisms (SNPs) and small insertions and deletions (INDELs), we applied a hybrid mode of DeepVariant^26^ that exploited both HiFi and Illumina read alignments^36^. Simultaneously, we used PEPPER-DeepVariant^27^ to generate additional SNP calls with ONT reads as it can yield high-quality SNP variants in difficult regions of the genome^36^ (**Supplementary Fig. 2**). We rigorously filtered all calls using Genotype Quality (GQ <= 30) and Variant Allele Frequency (VAF <= 0.5) to exclude any low-frequency false-positive calls, and filtered all of the suggested alternate corrections with Merfin^37^ to avoid introducing error *k*-mers (**Fig. 1b, Fig.3c))**. Finally, we ignored variants near the distal or proximal rDNA junctions on the short arms of the acrocentric chromosomes to avoid homogenizing the alleles from the un-assembled rDNAs. After merging all variant calls, we identified 993 small variants (<=50bp) that represented potential assembly errors and heterozygous sites. From these 993 assembly edits, about two-thirds were homopolymer corrections (512) or low-complexity micro-satellite repeats composed of 2 distinct bases in homopolymer-compressed space (hereby noted as “2-mer”) consistent with prior observations of HiFi sequence errors or bias^16^. Across all 617 loci, we evaluated the edit distribution using both Illumina and HiFi reads and found that the majority of Illumina reads supported the longer homopolymer or 2-mer repeat lengths compared to HiFi reads, thereby uncovering systemic biases in both homopolymer and 2-mer length in HiFi reads^16^ that caused the propagation of these errors into the consensus assembly sequence **(Fig.3d)**.

We used Parliament2^38^ and Sniffles^39^ to identify medium-sized (>50bp) assembly errors and heterozygous structural variants (SVs). To improve specificity, we only considered Sniffles calls supported by at least two long-read technologies (HiFi, ONT, and CLR) and Parliament2 calls supported by at least two SV callers. Similar to small variant detection, we excluded SVs called in the partial rDNA arrays and the HSat3 satellite repeat on chromosome 9. This pipeline identified a relatively small number of SV calls that we were able to manually curate via genome browsing. In total, we corrected three medium-sized assembly errors (replacing 1,998 bp of CHM13v0.9 sequence with 151 bp of new sequence) and we identified 44 heterozygous SVs (**Fig. 3a** and **Supplementary Fig. 3)**. We also identified a missing telomere sequence on the p-arm of chromosome 18 — a potential result of the string graph simplification process and confirmed through Bionano mapping (**Fig. 1b and 3b**). To correct this omission, we used the CHM13v0.9 graph to identify a set of HiFi reads expected to cover this locus^1^ and found ONT reads that mapped to the corresponding subtelomere and contained telomeric repeats. We used the ONT reads to derive a consensus chromosome 18 extension that was subsequently polished with the associated HiFi reads. After patching this telomere extension, we used Bionano alignments to confirm the accuracy of this locus (**Fig. 3b**). Altogether, the small and medium-sized variant calls along with the chromosome 18 telomere patch were combined into two distinct VCF files: a polishing edits file (homozygous ALT variants and the telomere patch) and a file for heterozygous variants (all other variants). We created the polished CHM13v1.0 assembly by incorporating these edits into the CHM13v0.9 with bcftools^40^.

**Figure 3.**
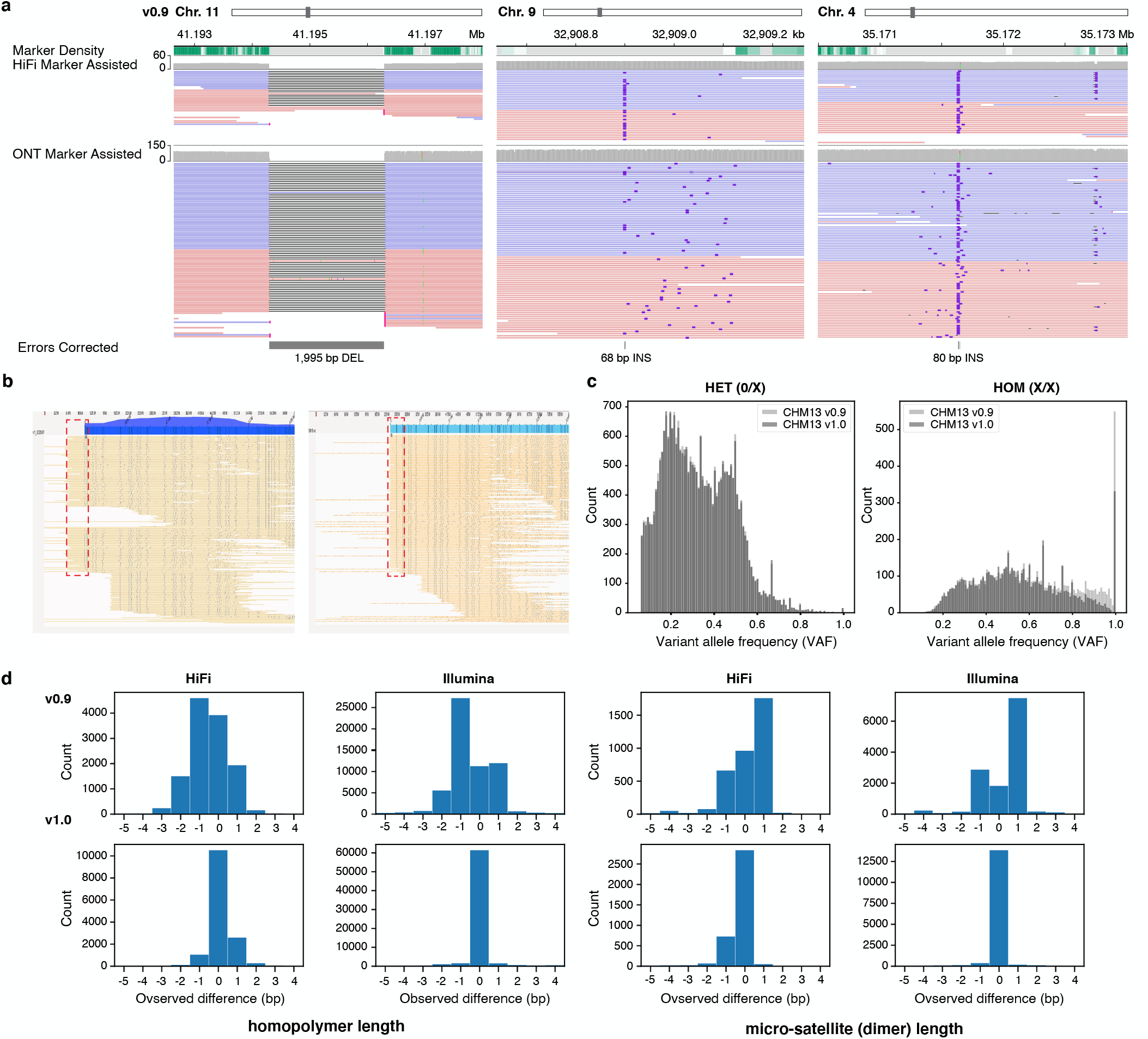
Errors corrected after polishing. **a**, Three corrected SV-like errors. **b**, Bionano optical maps indicating the missing telomeric sequence on Chr. 18 p-arm (left) with a higher than average mapping coverage. This excessive coverage was removed after adding the missing telomeric sequence (right) and most of the Bionano molecules end at the end of the sequence. **c**, Variant allele frequency (VAF) of each variant called by DeepVariant hybrid (HiFi + Illumina) mode, before and after polishing. Most of the high frequency variants (errors) are removed after polishing, which were called ‘Homozygous’ variants. **d**, Total number of reads in each observed length difference (bp) between the assembly and the aligned reads at each edit position. Positive numbers indicate more bases are found in the reads, while negative numbers indicate fewer bases in the reads. Both the homopolymer and micro-satellite (dimers in homopolymer compressed space) length difference became 0 after polishing.

We ensured polishing accuracy by extensive manual validation through visual inspection of the repeat-aware alignments, error *k*-mers, marker *k*-mers, and marker-assisted alignments. Here, we define “marker” *k*-mers as *k*-mers that occur only once in the assembly and in the expected single-copy coverage range of the read *k*-mer database and are highly likely to represent unique regions of the assembly (**Supplementary Fig. 4)**^41^. To generate marker-assisted alignments, we filtered Winnowmap2^34^ alignments to exclude any alignments that did not span marker *k*-mers (https://github.com/arangrhie/T2T-Polish/tree/master/marker_assisted). Our findings supported that most genomic loci contained a deep coverage of marker *k*-mers to facilitate marker-assisted alignment, except for a few highly repetitive regions (11.3 Mb in total) that lacked markers (termed “marker deserts”) (**Fig. 1a** and **Supplementary Fig. 4**). In parallel, we used TandemMapper^42^ to detect structural errors in all centromeric regions, including identified marker deserts. TandemMapper^42^ used locally unique markers for the detection of marker order and orientation discrepancies between the assembly and associated long reads. We manually validated all large polishing edits and heterozygous SVs, and many small loci were validated *ad hoc*.

### Validation of CHM13v1.0

Given the high completeness and accuracy standards of the T2T consortium, and knowing that polishing has the chance of introducing additional errors^37^, we took extra precautions to validate polishing edits and to ensure that edits did not degrade the quality of CHM13v0.9. First, we repeated self-alignment variant calling methods on CHM13v1.0, confirming that all edits made were correct (**Fig. 3a**). Through Bionano optical map alignments, we validated the structural accuracy of the chromosome 18 telomere patch and confirmed that all 46 telomeres were represented in CHM13v1.0 (**Fig. 3b**). Notably, our polishing led to a marked improvement in the distribution of GQ and VAF of small variant calls (**Fig. 3c and Supplementary Fig. 5a**). Our approach also increased the base level consensus accuracy from Q70.22 in CHM13v0.9 to Q72.62 in CHM13v1.0. Further, we found that error *k*-mers were uniformly distributed along each chromosome, suggesting that remaining errors were not clustered within certain genomic regions (**Supplementary Fig. 5b-c**). Upon re-evaluation of the homopolymers and 2-mers, we noted most of the biases we found in CHM13v0.9 from HiFi reads had been accurately removed, achieving an improved concordance with Illumina reads (**Fig. 3d**).

Overall, we made a total of 112 polishing edits (impacting 267 bp) in centromeric regions^43^, with 15 (35 bp) of these edits occurring specifically in centromeric alpha-satellite higher-order repeat arrays. We made 134 edits (4,975 bp) in non-satellite segmental duplications^2^. Moreover, the polishing edits were neither enriched nor depleted in satellite repeats and segmental duplications (p=0.85, permutation test), suggesting that non-masked repeats were not over- or under-corrected compared to the rest of the genome (**Supplementary Fig. 6**). Finally, through extensive manual inspection, we confirmed the reliability of the alignments for the three SV associated edits incorporated into CHM13v1.0 (**Supplementary Fig. 7**), and these efforts uncovered some heterozygous loci in the centromeres. These regions are under active investigation by the T2T consortium to both ensure their structure and understand their evolution^43^.

As an additional validation, we investigated potential rare or false collapses as well as rare or false duplications in CHM13v1.0. Here, based on *k*-mer estimates from both GRCh38 and CHM13v1.0 and from Illumina reads for 268 Simon’s Genome Diversity Project (SGDP) samples, we identified regions in CHM13v1.0 with a lower or higher copy number than both GRCh38 and 99% of the SGDP samples^2^. We found six regions of rare collapses in CHM13v1.0 that were not in GRCh38 (covering 205 kb, four from one single segmental duplication family). Both our HiFi read depth and Illumina *k*-mer-based copy number estimates suggest these six regions are likely rare copy number variants in CHM13 (e.g.,CHM13v1.0 has only a single copy of the 72 kb tandem duplication in GRCh38, **Fig. 4a)**. Additionally, we found that CHM13v1.0 had 33x fewer false or rare collapses than GRCh38 (∼185 loci (6.84 Mbp)) supporting erroneous missing copies in GRCh38^6^. We identified five regions (160 kb) with rare duplications in CHM13v1.0. This included a single 142 kb region that appeared to be a true, rare tandem duplication based on HiFi read depth and Illumina *k*-mer-based copy number estimates (**Fig. 4b**). Two of the smaller regions appeared to be true, rare tandem duplications, and two other small regions were identified during polishing as heterozygous or mosaic deletions, revealing potential tandem duplications arising during cell line division or immortalization. In summary, we found 7.5x fewer rare or falsely duplicated bases in CHM13v1.0 relative to the 12 likely falsely-duplicated regions affecting 1.2 Mb and 74 genes in GRCh38^6^, including the medically relevant genes: *CBS, CRYAA*, and *KCNE1*^*44*^.

**Figure 4.**
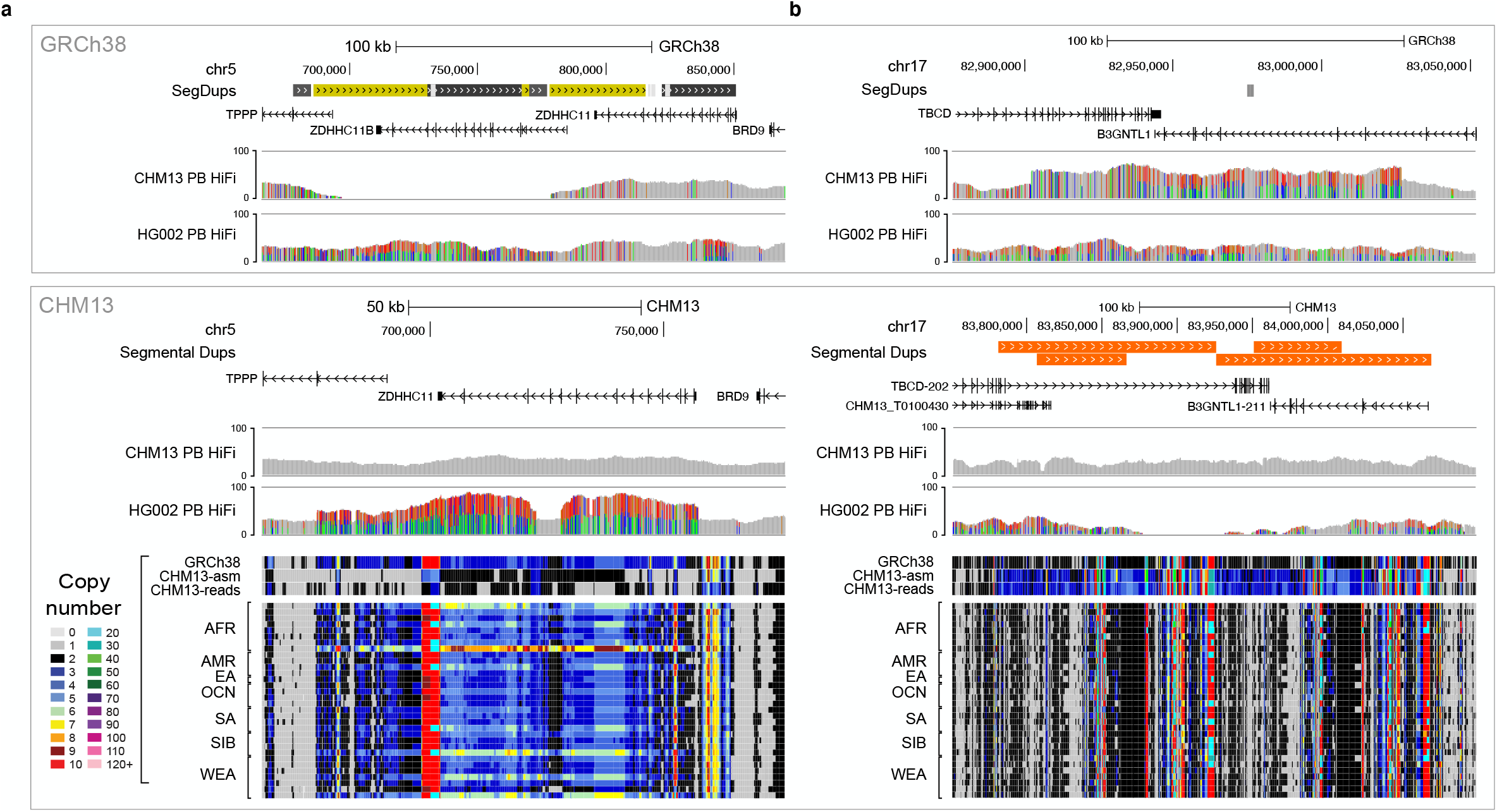
Examples of the largest CHM13 regions with a copy number in the reference that differs from GRCh38 and most individuals. **a**, One of the two largest examples of rare collapses in CHM13, where one copy of a common 72 kb tandem duplication is absent in CHM13. **b**, The largest rare duplication in CHM13, a 142 kb tandem duplication of sequence in GRCh38 that is rare in the population. CHM13 and HG002 PacBio HiFi coverage tracks are displayed for both references, GRCh38 (top) and CHM13v1.0 (bottom), to demonstrate that CHM13 reads support the CHM13 copy-number but HG002 reads are consistent with the GRCh38 copy-number. Read-depth copy-number estimates in CHM13 are shown at the bottom for ‘*k*-merized’ versions of GRCh38 and CHM13v1.0 references, CHM13 Illumina reads, and Illumina reads from a diverse subset (n=34) of SGDP individuals.

### Toward a completely polished sequence of a human genome

While evaluating CHM13v1.0, the T2T consortium successfully completed the construction of the rDNA models and their surrounding sequences on the *p*-arms of the five acrocentric chromosomes^1^. In parallel, we determined that all telomeric sequences remained unpolished. Specifically, in canonical [TTAGGG]n repeats, we found both HiFi read coverage dropouts and ONT strand bias impeded high quality variant calling **(Supplementary Fig. 8)**. For ONT, we observed only negative strands on the *p*-arm and only positive on the *q*-arm across all chromosomes; we suspect this is due to ONT reads never starting in telomeric sequences and only reading into them^41,45^. All existing variant callers for ONT reads require the presence of reads from both strands for accurate variant calling. Therefore, we tailored our PEPPER-based polishing approach and performed targeted telomere polishing to remove these errors remaining in telomeric sequences (**Methods**). Finally, automated polishing (described below), indicated that the *FAM156B* gene was heterozygous in CHM13v0.9 and CHM13v1.0 represented the rare minor allele (encoding a premature stop codon) at this locus. We replaced this minor allele with the other CHM13 allele encoding a full-length protein sequence. Overall, we made 454 telomere edits, producing longer stretches of maximum perfect matches to the canonical *k*-mer at each position across these telomeres compared to CHM13v1.0 (**Supplementary Fig. 9**). Combined with the parallel completion of the five rDNA arrays, our final round of polishing led to an improved QV of Q73.94 for CHM13v1.1.

Again, to ensure updates did not compromise the high accuracy of the assembly and to identify any remaining issues, we carried out an additional round of SV detection and manual curation using HiFi and ONT, classifying seven loci as remaining issues in CHM13v1.1 (**Supplementary Table 2**). Two loci located in the rDNA sequences appear to be a potential discrepancy between the model consensus sequence and actual reads or an artifact of mapping or sequencing bias. Lower consensus quality is indicated at two other loci, one detected with read alignments that were both low in coverage and identity, and one of which contained error *k*-mers detected by the hybrid dataset. One locus consisted of multiple insertions (<1kb) with breakpoints detected in low-complexity sequences associated with heterozygous variants and indicated a possible collapsed repeat (**Supplementary Fig. 10**) and an additional two loci joined and created an artificial chimeric haplotype (**Supplementary Fig. 11**). Additionally, we found 218 low coverage loci using HiFi (**Supplementary Table 3**), with 81.2% associated with GA-rich (78.0%). The remaining 41 loci had signatures of lower consensus quality and alignment identity, and 30 had error *k*-mers detected from the hybrid *k*-mer dataset. In contrast, we detected one low-coverage locus using ONT that overlapped the GA-rich model rDNA sequence. We associated most remaining loci, totalling only 544.8 kb or <0.02% of assembled sequence, with lower consensus quality in regions lacking unique markers. Overall, we found 394 heterozygous regions, including regions with clusters of heterozygous variants (https://github.com/mrvollger/nucfreq), totalling 317 sites (∼1.1 Mb).

We manually curated, both the breakpoints and alternate sequences associated with 47 heterozygous SVs, including sites previously inspected (CHM13v1.0) for SV-like error detection. We then investigated HiFi read alignment clippings and confirmed an association with clipping to both true heterozygous variant and spurious low frequency alignments. Additionally, we detected a further heterozygous inversion that went previously undetected.

### A comparison to automated assembly polishing

To demonstrate the efficacy of the customized DeepVariant-based approach, we compared our semi-automated polishing approach used to create CHM13v1.0 (Q72.62) to a popular state-of-the-art automated polishing tool, Racon^29,46^. To test, we iteratively polished CHM13v0.9 (three rounds) using Racon with PacBio HiFi alignments. While the QV improved from Q70.22 to Q70.48 after the first round of Racon polishing, it degraded with the subsequent second (Q70.26) and third (Q70.15) rounds, ultimately diminishing assembly accuracy. We also found that Racon incorporated 7,268 alternate alleles from heterozygous variants identified by DeepVariant, thus potentially causing undesirable haplotype-switching in haplotype-consistent blocks. To examine how Racon polished large, highly similar repetitive elements, we counted the number of corrections in non-overlapping 1 Mb windows of the CHM13v0.9 assembly and measured local polishing rates. Unlike CHM13v1.0, Racon polishing showed a clear right-tail in the distribution of polishing rates, indicating the presence of polishing “hotspots”, defined here as loci with >60 corrections/Mb **(Fig. 5a)**. The proximal and distal junctions of the rDNA units (masked from CHM13v1.0 polishing) were prevalent among these loci, a finding that reinforced the importance of masking rDNA loci to avoid overcorrection. We also found non-rDNA loci that were preferentially polished by Racon, including satellite repeats such as the highly repetitive HSat3 region in chromosome 9. Finally, CHM13v1.0 made two corrections, recovering as many protein-coding transcript’s open reading frames (ORFs), but Racon did not make these corrections. Racon also made 10 corrections that caused invalid ORFs in 30 transcripts (from nine genes) (**Fig. 5b**). Most of these corrections occurred at homopolymer repeats, consistent with our previous findings that homopolymer bias in HiFi reads could lead to false expansion or contraction of homopolymers during polishing.

**Figure 5.**
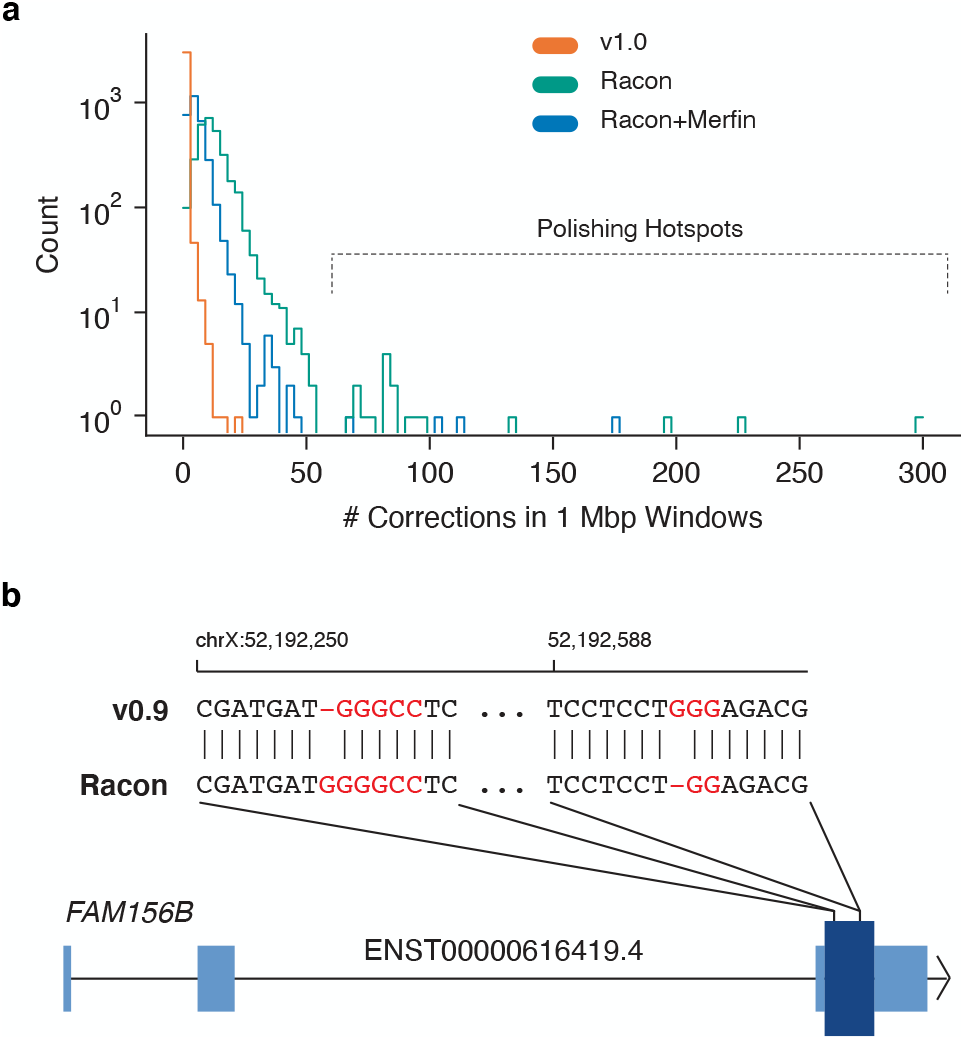
Errors made by automated polishing. **a**, The distribution of the number of polishing edits made in non-overlapping 1 Mb windows of the CHM13v0.9 assembly. **b**, Two Racon polishing edits causing false frameshift errors in the *FAM156B* gene. Light blue indicates UTR and dark blue indicates the single coding sequence exon. Highlighted sequence indicates GC-rich homopolymers.

To test if our newly automated *k*-mer based filtering tool could improve the specificity of automated polishing, we polished the CHM13v0.9 assembly with three rounds of Racon and Merfin (Racon+Merfin). After each round of polishing, Merfin filtered Racon edits that incorporated false assembly *k*-mers. As expected, the Racon+Merfin assembly QV monotonically increased from Q70.22 to Q77.34, Q77.99 and Q78.12. However, Racon+Merfin still incorporated 2,274 alternate alleles from heterozygous variants and polishing hotspots were still evident, suggesting that some repeats were overcorrected (**Fig. 5a**). Merfin mitigated the 10 ORF-invalidating Racon corrections, however, Merfin also failed to correct the two reading frame corrections made in CHM13v1.0 but not Racon. Overall, we suggest that Racon and Merfin can be used together as a highly effective automated polishing strategy for accurate draft assemblies. However, compared to these automated approaches, our custom polishing pipeline was superior for preserving haplotype consistency and avoiding repeat overcorrection.

## DISCUSSION

This work outlines our comprehensive polishing and evaluation strategy for the first complete human genome assembly using primarily PacBio HiFi for consensus sequence construction. Despite the high accuracy of both the data and the underlying graph, known errors existed that required polishing. Our polishing strategy required a deviation from the existing, aggressive, automated polishing tools and pipelines and a shift toward tailored evaluation and a “do no harm”, repeat aware, polishing pipeline that ensured its application did not compromise the high quality of the T2T Consortium’s CHM13v0.9. Pertinent to achieving an appropriate polishing strategy was our initial evaluation of CHM13v0.9, especially with respect to the complex repeats it uniquely revealed. This prior evaluation informed our dedicated effort to avoid overpolishing in unfinished areas such as the proximal and distal junctions of the rDNA arrays, large satellite repeat arrays, and regions subject to coverage dropouts. When evaluating HiFi and Illumina read homopolymer content we detected potential assembly errors or heterozygous variants that required polishing. Finally, we evaluated and confirmed the structural integrity of the CHM13v0.9 through Bionano, Strand-seq and Hi-C^1^. Through *k*-mer evaluations, we identified a drop-out of AT-rich *k*-mers in HiFi reads that could be recovered through Illumina reads through our evaluation of read *k*-mer content across sequencing technology platforms (35X HiFi, 100X Illumina PCR-free). Therefore, we created a merged *k*-mer database of both HiFi and Illumina reads to localise true errors to validate polishing edits.

Our evaluation of CHM13v0.9 identified coverage and sequencing platform biases that necessitated a custom and contextualised polishing model that capitalised on the wealth of available data and exploited the advantages of each sequencing platform to call both small SNVs and medium-sized SVs. Here, we developed an approach to increase variant calling specificity by using multiple SV and SNV calling tools^26,38,39^, one specifically designed to conservatively address the centromeric regions^42^. We followed by merging called variants to produce the final call set and filtering to prevent the introduction of erroneous *k*-mers^37^. To identify potential false-positive calls, we extensively validated both small and large variant calls through manual inspection of both self- and marker-assisted alignments. Here, we developed specific methods for telomere end polishing to cater for sequence platform biases causing natural decreases in coverage. We found that due the HiFi coverage dropouts that ONT reads were the only reliable data source when polishing these long telomeric repeats; however, strand bias needed to be accommodated when polishing^45^. As existing variant callers avoid unmappable regions caused by coverage dropouts and rely heavily on support from both strands for accurate variant calling, we developed dedicated methods to accurately polish these regions. Overall, our customised and context-specific strategy carefully navigated the identified idiosyncrasies of CHM13v0.9 and called for just 1,457 corrections (Q73.9) including: *p*-arm of chromosome 18; 454 telomere corrections; 1 large deletion; 2 large insertions; 993 SNPs; 113 small insertions and 880 small deletions. Although the final CHM13v1.1 is highly accurate, we identified 225 loci that were recalcitrant to validation, and we have documented these loci along with 394 heterozygous loci (317 merged loci) (https://github.com/marbl/CHM13-issues/).

The high accuracy of CHM13v1.1 showcases the effectiveness of our informed selection and implementation of appropriate repeat-aware aligners^34,42^, *k*-mer evaluation and filtration tools, and highly accurate and sensitive variant callers^27,37^ whilst also highlighting the utility of capitalising on the synergistic nature of multiple sequencing technology platforms. The minimal number of corrections implemented by our approach and uniform coverage (99.86%) exemplifies the high accuracy of the initial graph construction, with sequencing biases being associated with the remaining coverage fluctuations (223 regions were regions of HiFi dropouts, 77.5% found in GA/TC-rich, and AT-rich satellite sequences such as HSat2/3 and HSat1, were associated with in HiFi coverage increases and ONT coverage depletion, respectively).

Achieving a complete human genome sequence was made possible by several factors including, the low level of heterozygosity of the CHM13 cell line, recent advancements in sequencing technologies, customised algorithms for string graphs to better resolve repeats, and a dedicated team for manual validation - a result not yet to be expected by current automated genome assembly algorithms^16,17^. Moreover, current automated polishing and variant calling tools are limited in their ability to stringently polish or indeed distinguish error from true heterozygous variants on genome assemblies of >Q70 where the majority of the consensus sequence is already haplotype-consistent, nor have they been fine tuned to properly correct errors caused by sequence bias. Therefore, if implemented without manual curation they could lead to false positives, particularly in highly repetitive regions such as rDNA, centromeric satellites and segmental duplications. However, despite the unique, unconventional, semi-automated nature of our polishing and evaluation endeavor, recent trends in DNA sequencing and genome assembly algorithms suggest that CHM13v1.1 is just a preview of an imminent wave of high-quality T2T reference genomes in other species^47–49^. It is therefore critical our lessons outlined be incorporated into the next generation of automated bioinformatics tools^29,34,42,37^ and to ensure that as our “norm” for genome assembly quality transforms, a simultaneous and concerted transformation of our evaluation and polishing methods and their automation is pursued.

Complete and highly accurate genome assemblies will become more routine in the next several years, exposing the limitations of many existing polishing and validation tools. Implementing automated polishing processes alone runs the risk of introducing more errors than fixes in genomes of this quality, and this is why our “do no harm”, semi-manual, approach was necessary for the completion of the first human genome. However, the use of phased reads to mitigate heterozygous switches during automated polishing followed by subsequent automated *k*-mer based filtration of the variants called has the potential to limit harm and achieve a genome assembly of a quality sufficient for most assembly projects. However, caution should be exercised specifically in the homopolymer and microsatellite regions prone to sequencing biases causing repeat shortening. Moreover, multiple data types should be used to inform polishing strategies in these regions. Here, we present the tools and methods we developed to polish the first complete human genome; however, the lessons learned extend far beyond the scope of this milestone, and will be implemented and further developed by ongoing efforts such as the Human Pangenome Reference Consortium^50^ and Vertebrate Genomes Project^22^. To maximise their utility, the future requires automation of these tools to accurately map the previously unmappable through repeat aware alignment; streamline assembly visualisation for curation of variant calls; filter variant calls through *k*-mer based evaluation; and sensitively and reliably identify variants and errors from telomere to telomere.

## ACKNOWLEDGEMENTS

This work was supported by the Intramural Research Program of the National Human Genome Research Institute (NHGRI), National Institutes of Health (NIH) (AM, CJ, SK, AMP, AR); National Science Foundation: DBI-1350041 and IOS-1732253 (MA); NIH/NHGRI R01HG010485, U41HG010972, U01HG010961, U24HG011853, OT2OD026682 (KS, BP); HHMI (GF); Wellcome WT206194 (KH); NIGMS F32 GM134558 (GAL); NIH/NHGRI R01 1R01HG011274-01, NIH/NHGRI R21 1R21HG010548-01, and NIH/NHGRI U01 1U01HG010971 (KM) ; St. Petersburg State University grant ID PURE 73023573 (AM); NIH/NHGRI R01 HG006677 (AS); Fulbright Fellowship (DCS); Wellcome WT206194 (JMW); Intramural funding at the National Institute of Standards and Technology (JZ). This work utilized the computational resources of the NIH HPC Biowulf cluster (https://hpc.nih.gov). Certain commercial equipment, instruments, or materials are identified to specify adequately experimental conditions or reported results. Such identification does not imply recommendation or endorsement by the National Institute of Standards and Technology, nor does it imply that the equipment, instruments, or materials identified are necessarily the best available for the purpose.

## AUTHOR CONTRIBUTION

AR and AMP conceived and supervised the project. AMM, KS, GF, KH, JMDW, and AR performed the pre-polishing evaluation. KS, MA, AVB, AF, CJ, AM, BP, and AR aligned reads and called variants. AMM, KS, MA, GF, AF, KHM, AM, JMZ, and AR manually validated variant calls. DCS and JMZ performed the gene collapse and expansion analysis. KS, MA, AVB, GAL, KHM, AM, and AR identified and curated heterozygous and “issues” loci. KS, MA, SK, and BP patched and polished the telomeres. AMM, MA, AS, and IS performed automated polishing. AMM, KS, MA and AR wrote the manuscript, with assistance from all authors. All authors approved of the final manuscript.

## ONLINE METHODS

Codes used in this manuscript are openly available on our GitHub (https://github.com/arangrhie/T2T-Polish) unless otherwise noted.

### Evaluating homopolymer concordance

By analyzing the homopolymer length agreement, we assessed sequencing platform-specific biases between reads and the assembly using both Illumina and HiFi reads through the runLengthMatrix submodule of Margin (https://github.com/UCSC-nanopore-cgl/margin). Here, we used Margin to convert the assembly sequence to a run-length encoded (RLE) sequence. For example, the sequence ACTTG became (ACTG, {1,1,2,1}) where ACTG represented the encoded sequence, and {1,1,2,1} represented the run-length for each nucleotide base. While encoding the sequence to run-length, Margin created a map of positions in the assembly to the RLE position. Using the position map, Margin converted the raw sequence alignment to run-length alignment by iterating through the matches between the read and the assembly and keeping track of the previous match in RLE space. This way, Margin created a matrix where each row represents a run-length of a nucleotide base observed in the reads, and each column represents the run-length observed at the corresponding position in the assembly where the read mapped.

### Identifying potential polishing edits and heterozygous variants

To find potential polishing edits and heterozygous variants, we aligned a variety of public CHM13 WGS sequencing reads to CHM13v0.9 (https://github.com/marbl/CHM13). We refer to these alignments as “self-alignments” as both the query reads and reference assembly represent the CHM13 genome. Further, we aligned Illumina reads with BWA-MEM (v0.7.15)^51^ and removed PCR duplicate-like redundancies using ‘biobambam2 bamsormadup’ (v2.0.87)^52^ with default parameters. Pacific Biosciences Continuous Long Read (CLR) and Circular Consensus Sequencing (CCS/HiFi), and Oxford Nanopore (ONT) reads were aligned using Winnowmap2 (v1.1).

We used both Illumina and HiFi read alignments to call SNPs and indels with the “hybrid” model of DeepVariant (v1.0) but only ONT alignments were used to call SNPs using PEPPER-DeepVariant (v1.0)^27^. To exclude potentially spurious variant calls, we removed variants with low allele fraction support or low genotype quality (VAF<=0.5, GQ<=30 for Illumina/HiFi, and GQ<=25 for ONT). We then combined Illumina/HiFi hybrid and ONT variant calls using a custom script (https://github.com/kishwarshafin/T2T_polishing_scripts/blob/master/polishing_merge_script/vcf_merge_t2t.py). Finally, we filtered small polishing edits using Merfin^37^ to ensure all retained edits did not introduce any false 21-mers that were absent from the Illumina or HiFi reads.

Our approach implemented structural variant (SV) inference tools to detect medium-sized polishing edits and structural heterozygosity. For short-read-based SV calling, we used Illumina alignments as input to Parliament2^38^ using default settings. For long-read SV calling, we relied on HiFi, CLR, and ONT alignments to call SVs with Sniffles^39^ (v1.0.12, -s 3 -d 500 -n -1) and we removed all SVs with less than 30% of reads supporting the ALT allele. After this, we generated and refined insertion and deletion sequences with Iris (v1.0.3, using Minimap2^53^ and Racon^29^ for aligning and polishing, respectively)(https://github.com/mkirsche/Iris). Our approach yielded three independent technology-specific call sets that we merged using Jasmine (v1.0.2, max_dist=500 min_seq_id=0.3 spec_reads=3 --output_genotypes)^54^. Through manual inspection in IGV we validated all long-read variant calls longer than 30 bp supported by at least two technologies and all short-read SV calls^55^.

Our approach combined small and structural variant calls into two distinct VCF files: one for potential polishing edits (homozygous ALT alleles) and one for putative heterozygous variants (heterozygous ALT alleles) and we excluded all edits within known problematic loci - prone to producing false variant calls (rDNA gaps as well as the large HSat3 region on chromosome 9). To generate the CHM13v1.0, we applied ‘bcftools consensus’ (v1.10.2-140-gc40d090) to incorporate the suggested polishing edits into CHM13v0.9^56^ and repeated same previously detailed methods with respect to CHM13v1.0 to ensure that no additional polishing edits were apparent and to call heterozygous loci.

### Patching the chromosome 18 p-arm telomere

As a result of the string graph simplification process, we found a telomere missing from the graph representing the *p*-arm of chromosome 18. We identified five ONT reads associated with these telomeric sequences using the telomere pipeline developed by the VGP (https://github.com/VGP/vgp-assembly). Using these reads we ran Medaka (v1.0.3)^28^ to generate a consensus sequence and manually patched it into the assembly [https://github.com/malonge/PatchPolish]. We obtained seven matching HiFi reads, not in the assembly graph and confirmed to have telomeric repeats, and used Racon^29^ to further polish. In total, we added 4,862 bp of telomere sequence to the start of chromosome 18.

### Evaluating polishing accuracy

We repeated self-alignment variant calling methods on CHM13v1.0 and confirmed that no additional polishing errors were apparent. In addition to the self-alignments used for polishing and heterozygous variant calling, we derived marker-assisted alignments from previously created HiFi, CLR, and ONT Winnowmap2 alignments^35^. For marker-assisted alignment production, we removed Winnowmap2 alignments that did not span “marker” *k*-mers. We define marker *k*-mers as any 21-mer present once in CHM13v1.0 and between 42 and 133 times in the Illumina reads^41^ and filtered reads using technology specific length thresholds with HiFi having a 10kbp, CLR a 1kbp and ONT a 25kbp threshold. Our approach relied on both CHM13v1.0 self-alignments and marker-assisted alignments for manual inspection.

We also assessed the genome assembly using Merqury QV estimations based on 21-mer databases we created for both Illumina PCR-free and HiFi reads^32^. Following this, we derived a “hybrid” Merqury *k*-mer database using Meryl by combining Illumina and HiFi *k*-mers that occured over 23 and 4 times, respectively. To match the *k*-mer frequency in each copy number, we increased *k*-mer frequency in HiFi reads by 4 and we divided Illumina *k*-mer frequency by 3 and combined the *k*-mer databases by taking the union of the maximum frequency. To identify regions with rare collapses or rare duplications in CHM13v1.0, we compared copy number estimates of CHM13v1.0 to copy number estimates of 268 human genomes (Simons Genome Diversity Project; SGDP) using short reads. We averaged these copy number estimates for each genome across 1 kbp windows and we flagged a potential false or rare duplication if the copy number in CHM13v1.0 was greater than the copy number in 99% of the other genomes and GRCh38. Moreover, we flagged a potential false or rare collapse if the copy number in CHM13v1.0 was less than the copy number in 99% of the other genomes and GRCh38 and assigned all flagged regions a value of 1 and unflagged regions a value of 0. We included GRCh38 in this analysis to help remove rare technical artifacts where the assembly-based *k*-mer copy number estimate is systematically different from the Illumina read-based *k*-mer estimate. To filter the flagged regions, we used a median filter approach with a window size of 3 kbp where the binary value of each 1 kbp region was replaced with the median value of the complete window. Finally, we merged all adjacent flagged regions and reported the start and end coordinates with respect to CHM13v1.0, and we curated and removed flagged regions if they overlapped LINEs as SGDP copy number estimates are less reliable in these high copy number repeats.

### Polishing enrichment or depletion within repeats

We performed a permutation test to check if our polishing pipeline suggested significantly more or fewer polishing edits within repeats compared to the rest of the genome. We established two distinct samples of genomic intervals. For the first, we randomly sampled 20,000 100 kbp windows from the genome and removed any windows that intersected repeats. For the second, we randomly sampled 20,000 100 kbp windows and removed any windows that intersected non-repeats. By measuring the number of polishing edits in each 100 kbp window, we established two different random distributions of polishing rates: one within and one without repeats. We utilised SciPy *stats.ttest_ind* using 10,000 permutations to derive our p-value^57^.

### Telomere polishing

We employed a targeted polishing of telomeres by retraining PEPPER on HG002 chr20 with all forward strand reads removed to correct for the original model’s dependence on having reads from both strands. Using this retrained model, we generated a set of candidate variants in the telomere regions and the coverage depth was calculated using samtools depth. Finally, we implemented a custom script (https://github.com/kishwarshafin/T2T_polishing_scripts/blob/master/telomere_variants/generate_telomere_edits.py) that took these candidate variants and calculated the Levenshtein distance between the canonical telomere *k*-mer and the sequence we derived after the candidate variant had been applied. We selected only those variants as true telomere edits if the candidate had a minimum allele frequency of 0.5, a minimum genotype quality of 2 and reduced the Levenshtein distance to the canonical telomere *k*-mer when compared to the existing telomere sequence. Further, we trimmed the consensus sequence where ONT read depth support was lower than 5.

We employed SV detection to identify regions with low coverage support, excessive read clippings, and enriched secondary alleles, and to further ensure that accuracy was not compromised but also to identify and document outstanding issues with CHM13v1.1 (**Fig. 1a**). On inspection of both Winnowmap2^34^ and Minimap2^53^ read clippings, artificial alignment breaks were highlighted that caused clipping and coverage drops in regions with highly identical satellite sequences. Notably, we did not identify these breaks in alignments from TandemMapper^42^, a more conservative aligner specifically designed for alignment in satellite repeats. On further inspection of clipped reads, we found the chaining algorithm of Winnowmap2 handled lower confidence alignment blocks incorrectly, and so we updated accordingly (v2.0 to v2.01) for all future evaluations of both CHM13v1.0 and CHM13v1.1 (**Supplementary Fig. 12)**.

### Comparison to automated polishing approaches

To evaluate our newly proposed approach to polishing, we compared it to the off-the-shelf tools available for HiFi reads. We performed three rounds of iterative polishing using the Racon consensus tool with each iteration including the following steps. (1) Alignment of input HiFi reads to the input target sequences using Winnowmap 1.11 (https://github.com/marbl/Winnowmap/releases/tag/v1.11; options: “--MD -W bad_mers.txt -ax map-pb”). We used CHM13v0.9 (unpolished) as the first iteration target, while every following iteration used the polished output of the previous stage as the input target. (2) We filtered secondary alignments and alignments with excessive clipping using the “falconc bam-filter-clipped” tool (available in the “pbipa” Bioconda package; options: “falconc bam-filter-clipped -t -F 0×104”). By default, maximum clipping on either left or right side of an alignment is set to 100bp, but this was applied only if the alignment was located at least 25bp from the target sequence end (to prevent clipping due to contig which could otherwise cause false alignment filtering). (3) Finally, we used Racon (https://github.com/isovic/racon, branch “liftover”, commit: 73e4311) to polish the target sequences using these filtered alignments. For the purposes of this work, we extended the “master” branch of Racon to include two custom features: BED selection of regions for polishing and logging all changes introduced to the input draft assembly to produce the final polished output (in VCF, PAF or SAM format). We then ran Racon with default options with the exception of two new logging options: “-L out_prefix -S” implemented to store the liftover information between the input and output sequences. We used Liftoff (v1.6.0, -chroms -copies -exclude_partial -polish) using gencode v35 to annotate each of the polished assemblies^58,59^.

## Notes

### Competing Interest Statement

Ivan Sovic is employed by Pacific BioSciences Inc.

