## Supplementary_Figures&Tables for "Chasing perfection: validation and polishing strategies for telomere-to-telomere genome assemblies"

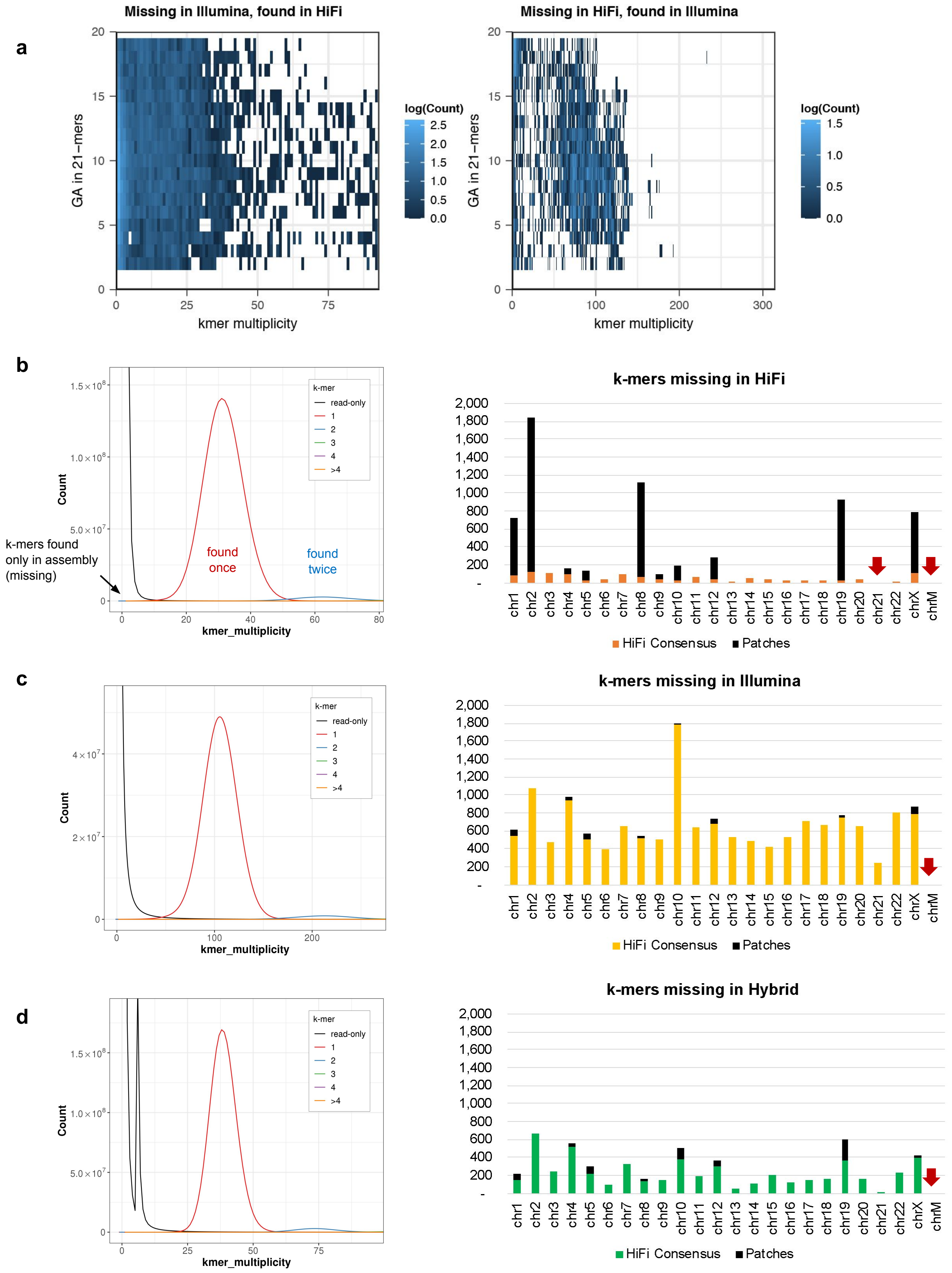


**Supplementary Fig. 1 | Sequencing biases observed in missing kmers. a,** missing k-mers with its GA composition. **b-d**, v0.9 assembly and k-mer copy number spectrum from HiFi, Illumina, and hybrid k-mer sets (left) and per-chromosome missing (likely error) k-mer counts from the HiFi derived consensus or patches (right). Most missing k-mers in HiFi overlapped sequences from patched regions. No missing k-mer was found on Chromosomes indicated with red arrows.

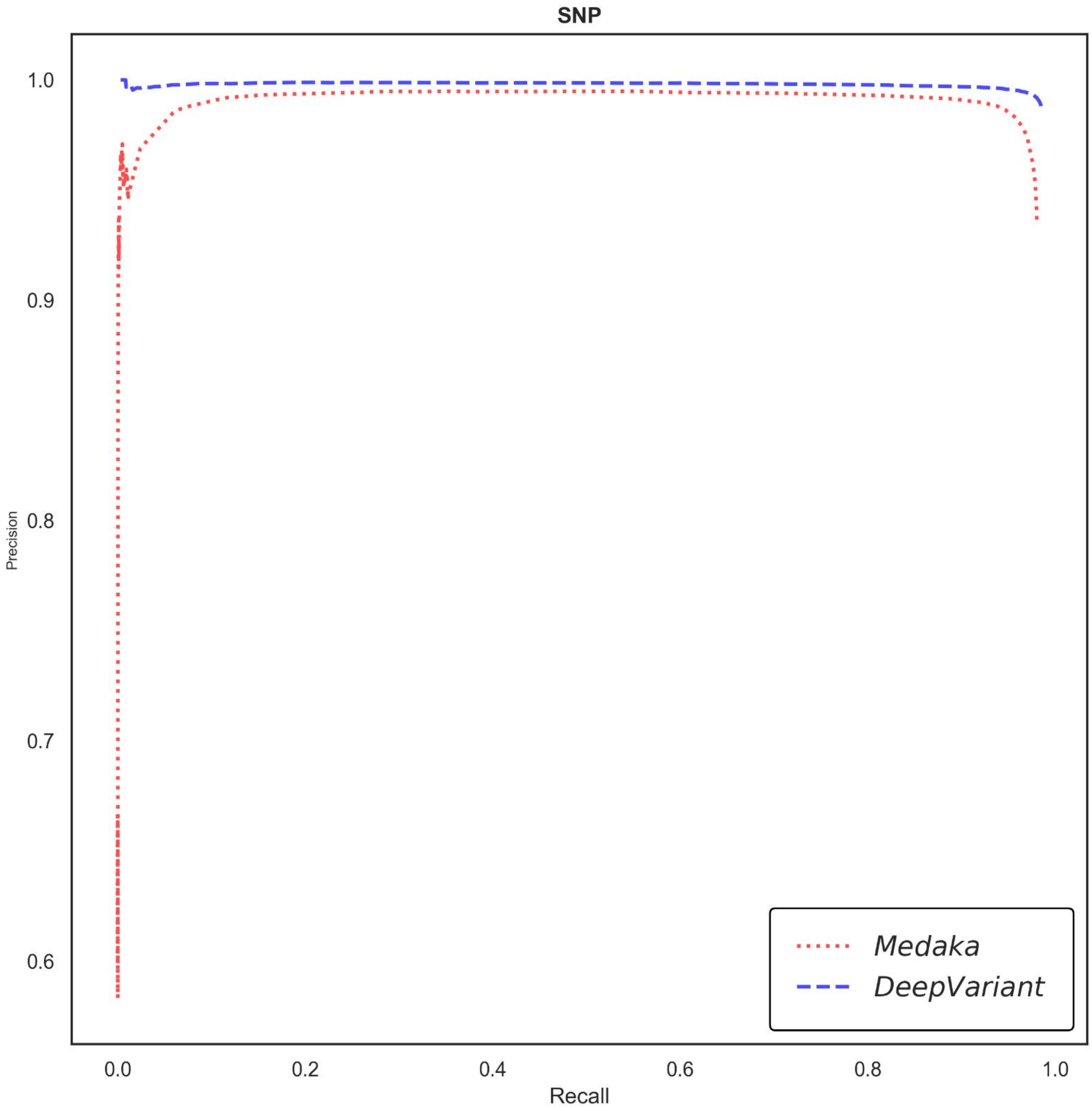


**Supplementary Fig. 2 | SNV-like error filtering.** ONT PEPPER-DeepVariant SNP call were more reliable with both higher precision and recall over Medaka.

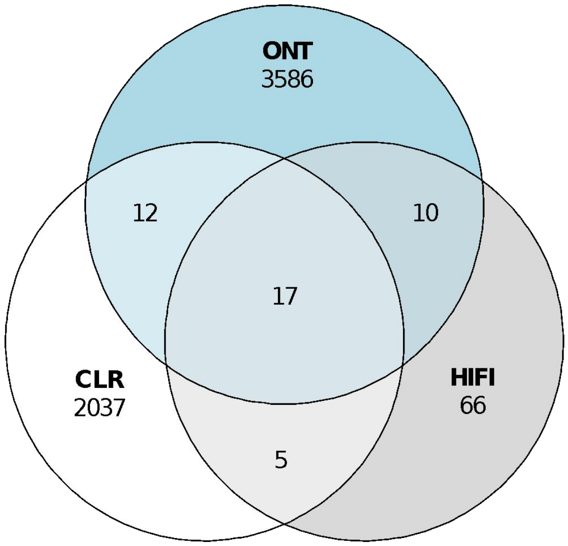


**Supplementary Fig. 3 | Number of SV-like errors called from long-read platforms.**

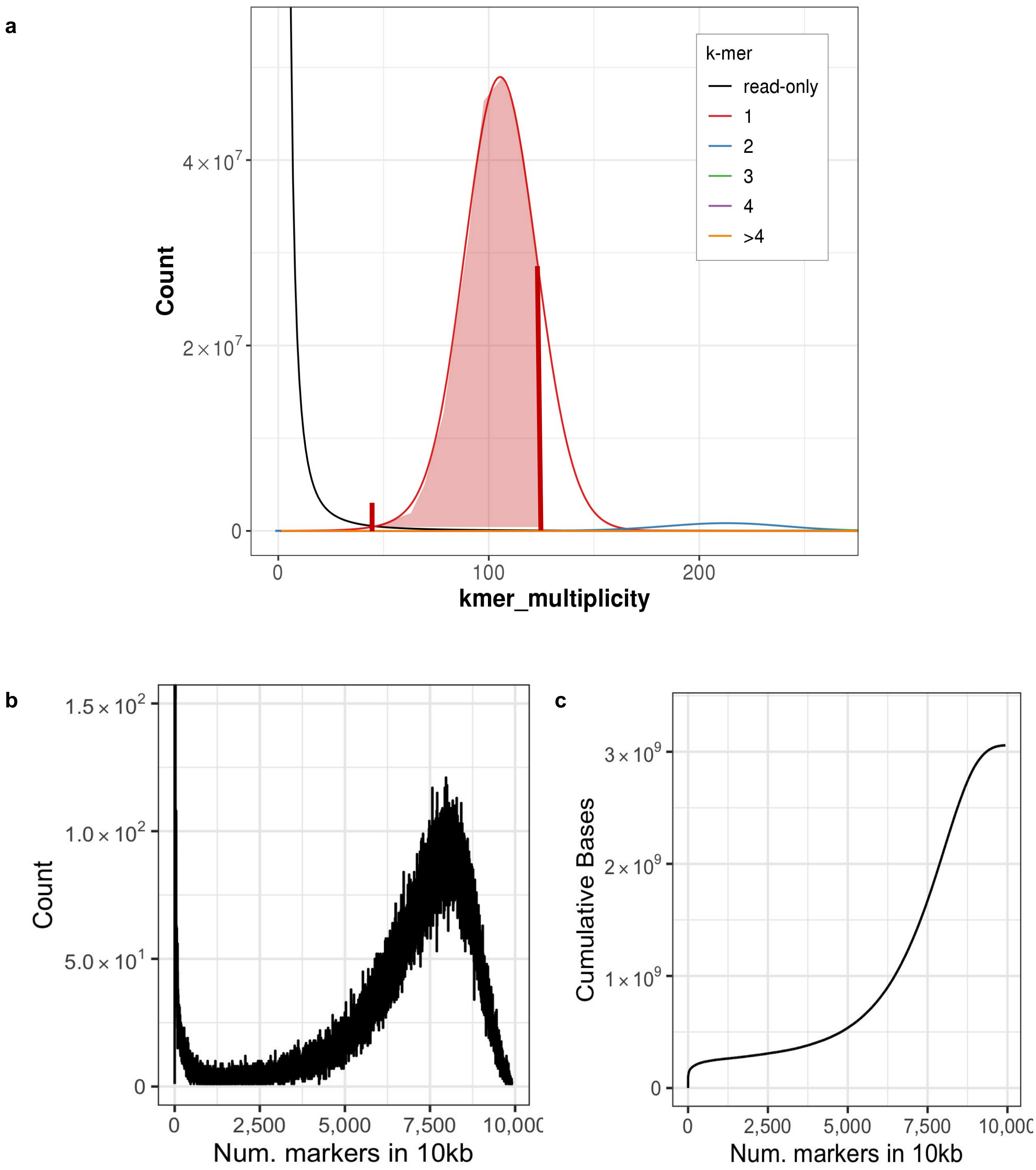


**Supplementary Fig. 4. | Globally unique single-copy kmers used for marker assisted alignment. a.** Range of k-mer counts defined as ‘single-copy’ markers from Illumina reads and in the assembly. The cutoffs were chosen to minimize inclusion of low-frequency erroneous kmers and 2-copy k-mers. **b**. Number of markers in every 10 kb window. **c**. Cumulative number of bases covered by the number of markers in each 10 kb window.

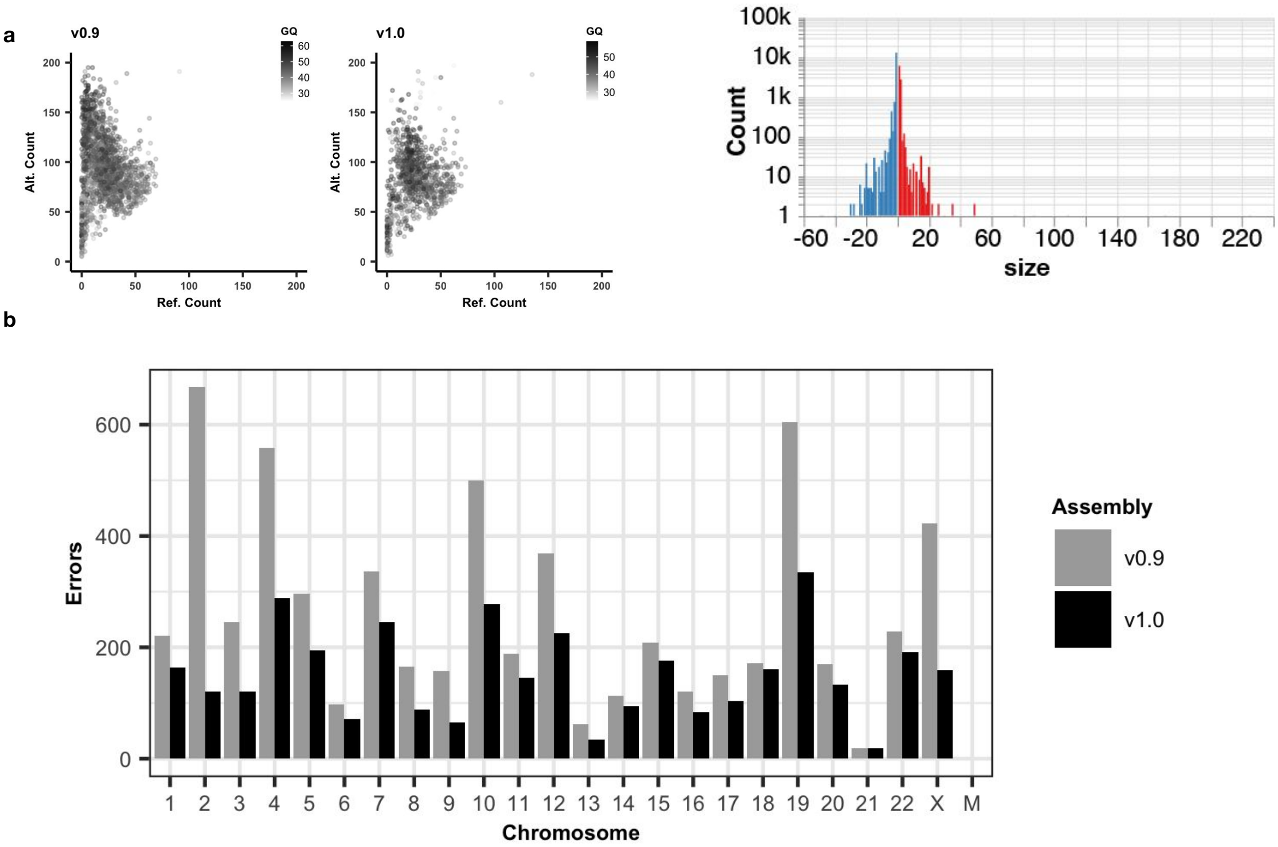


**Supplementary Fig. 5 | Post-polishing evaluation. a.** Left, genotype quality and number of reads supporting the reference and alternate alleles from the combined Illumina-hifi hybrid and ONT homozygous variant calls, with AF > 0.5. Right, balanced insertion (red) and deletion (blue) length distribution from the Illumina-HiFi hybrid DeepVariant heterozygous calls in v1.0. **b**. Number of errors detected in each chromosome, before and after polishing.

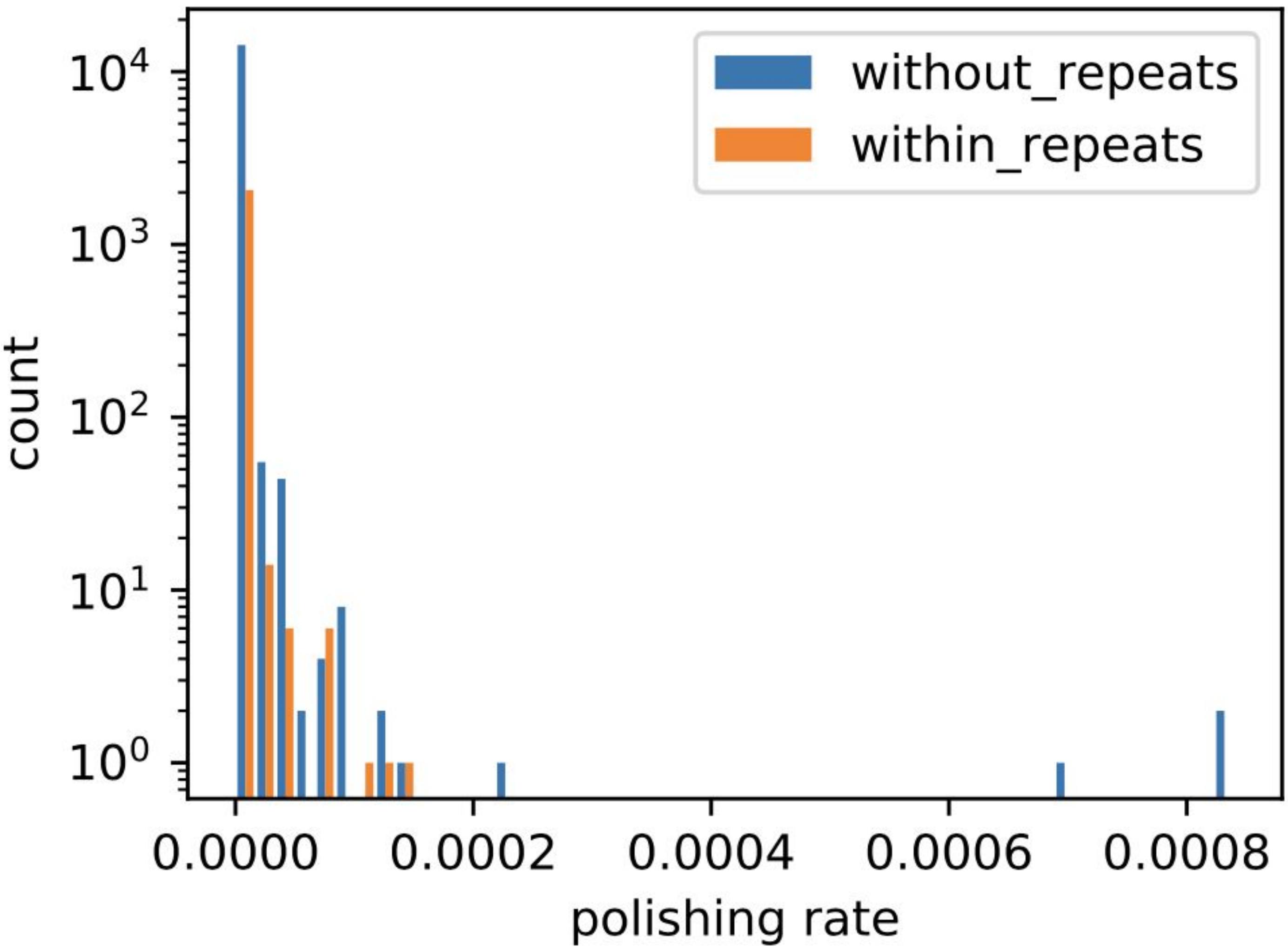


**Supplementary Fig. 6 | Polishing inside and outside of repeats.** The distribution of v0.9 polishing rates within and without repeats.

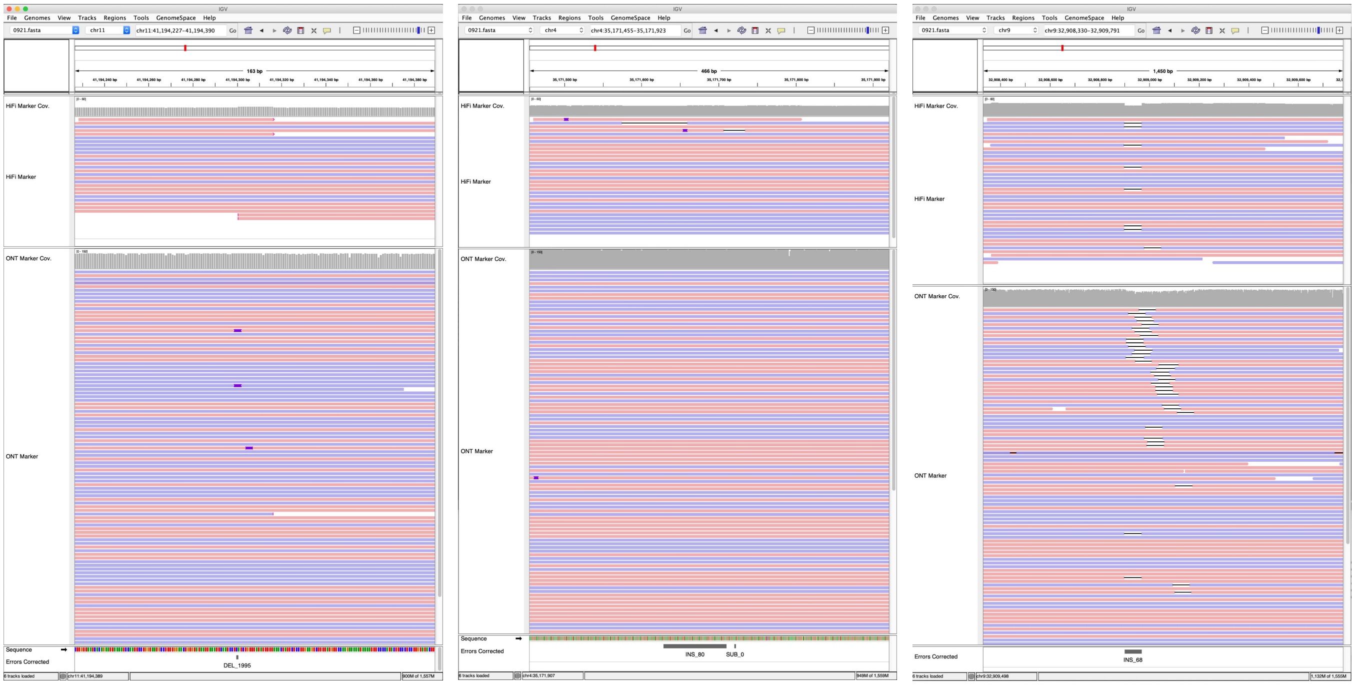


**Supplementary Fig. 7 | Three SV-like errors corrected.** HiFi and ONT marker assisted alignments, post correction of the 3 large SV-like edits visualized with IGV. HiFi coverage track is shown in data range up to 60, ONT up to 150. Clipped reads are flagged for >100bp. INDELs smaller than 10bp are not shown. Reads are colored by strands; positive in red and negative in blue.

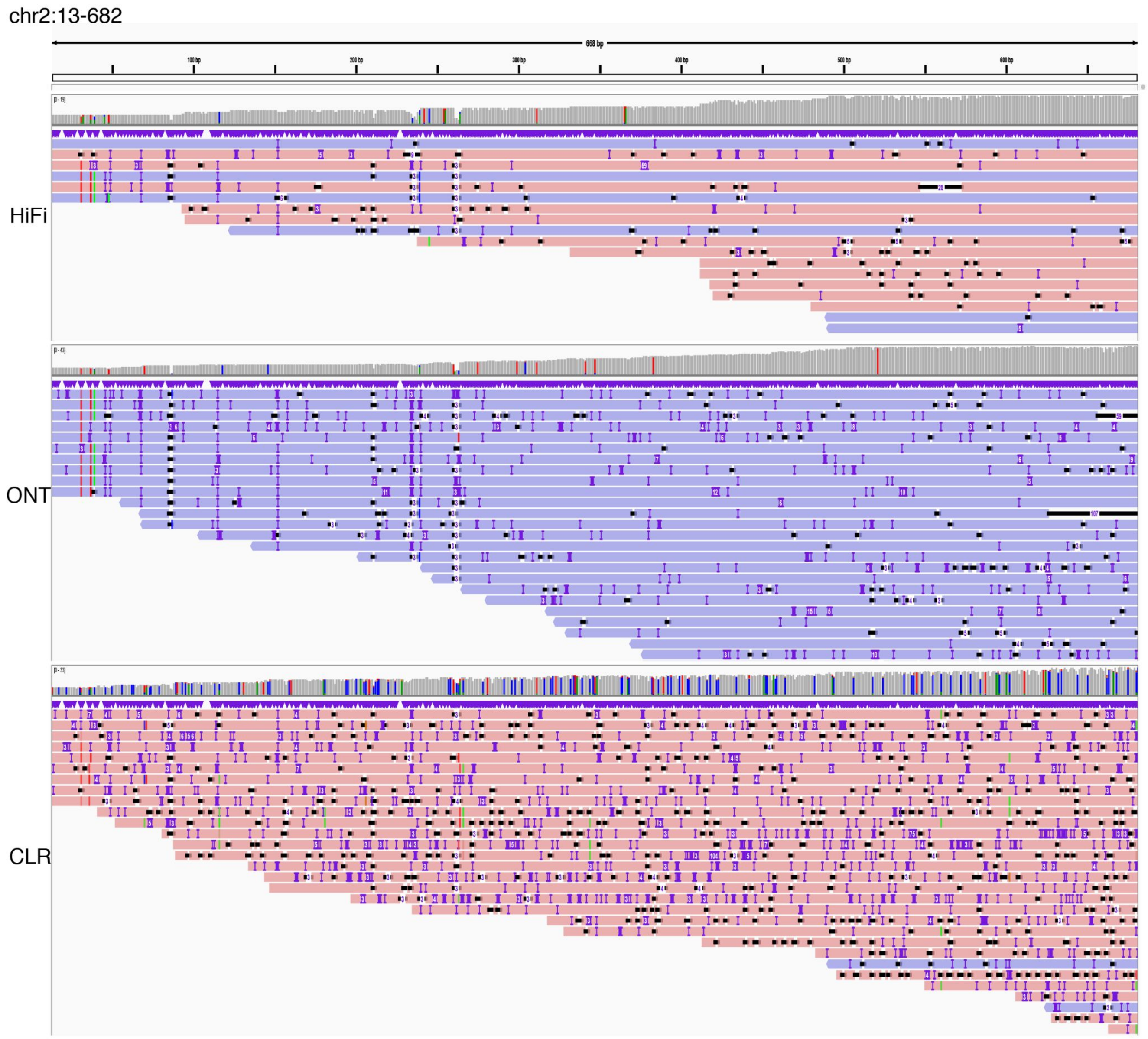


**Supplementary Fig. 8 | An illustration of chr 2 telomere sequence reads from HiFi, ONT and CLR platform.**

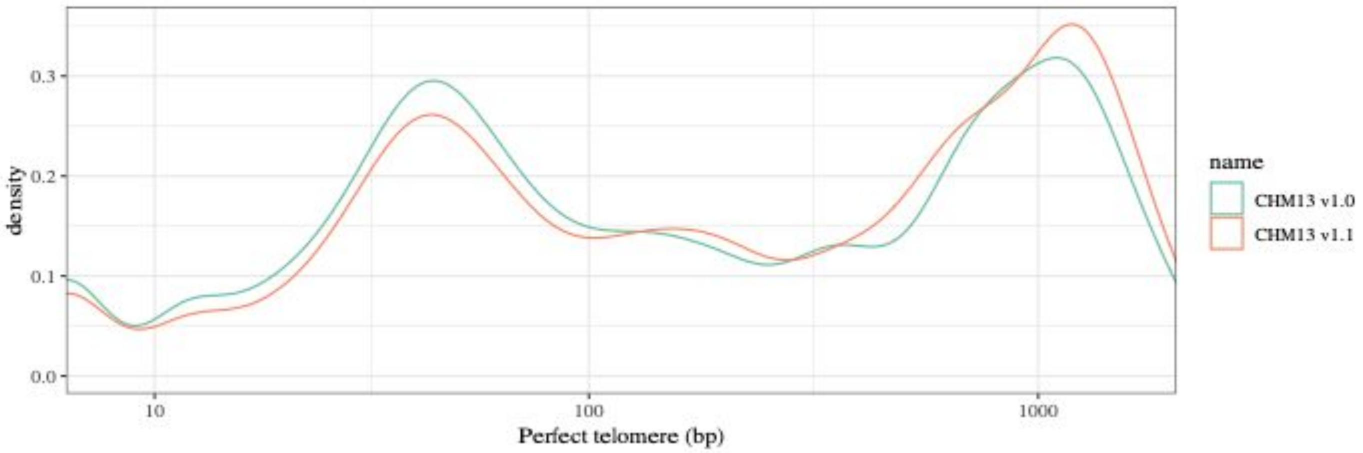


**Supplementary Fig. 9 | Distribution of maximum perfect match to the canonical k-mer observed at each position in the telomere before (CHM13 v1.0) and after (CHM13 v1.1) polishing the telomeres.**

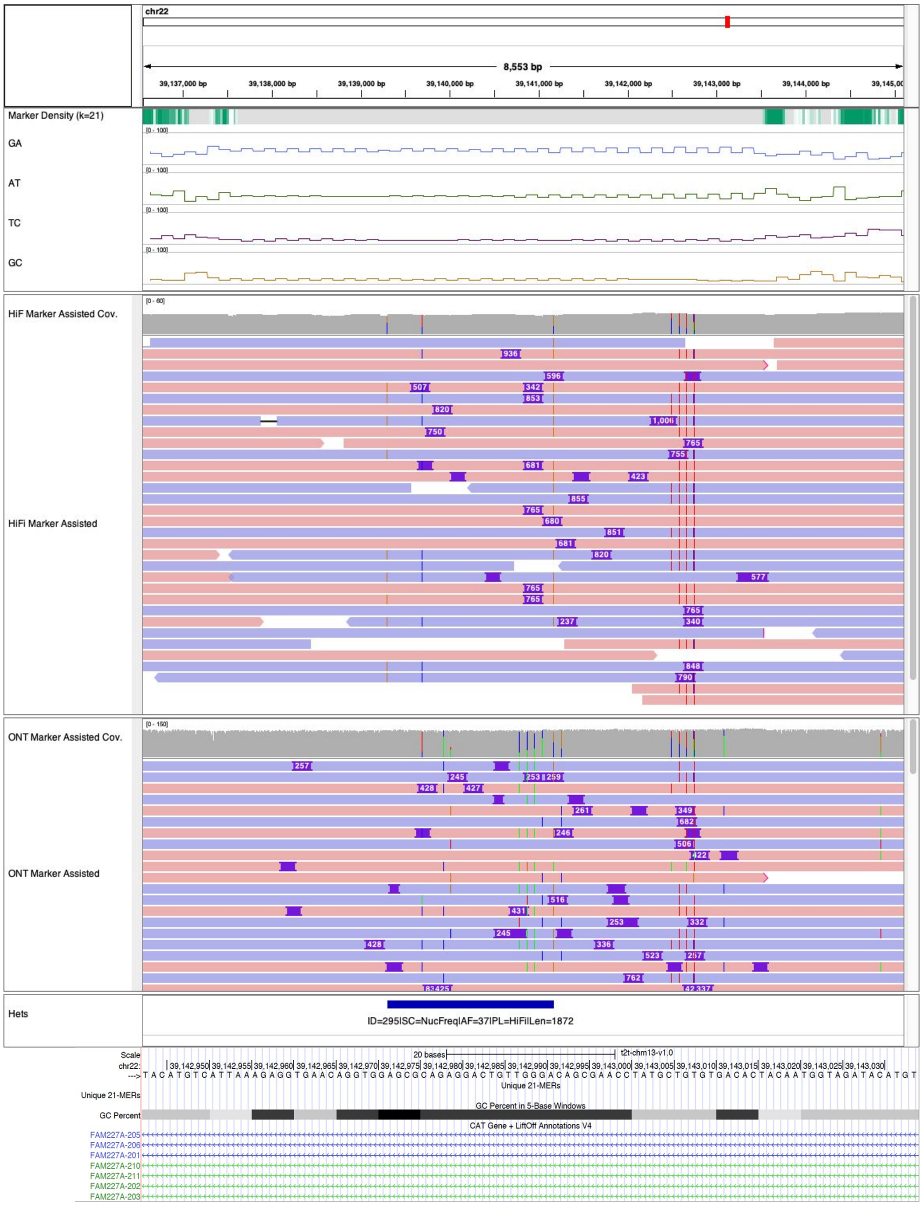


**Supplementary Fig. 10 | Collapsed simple tandem repeat.** The collapse in the Intronic sequences of gene FAM227A was undetected, due to the variable insertion breakpoints and insertion length in the HiFi and ONT alignments. The panels above the alignments show marker density and percent microsatellites (GA / AT / TC / GC) in each 64 bp window, which indicates this region is highly repetitive with GA enriched sequences, which later alternates with AT enriched sequences.

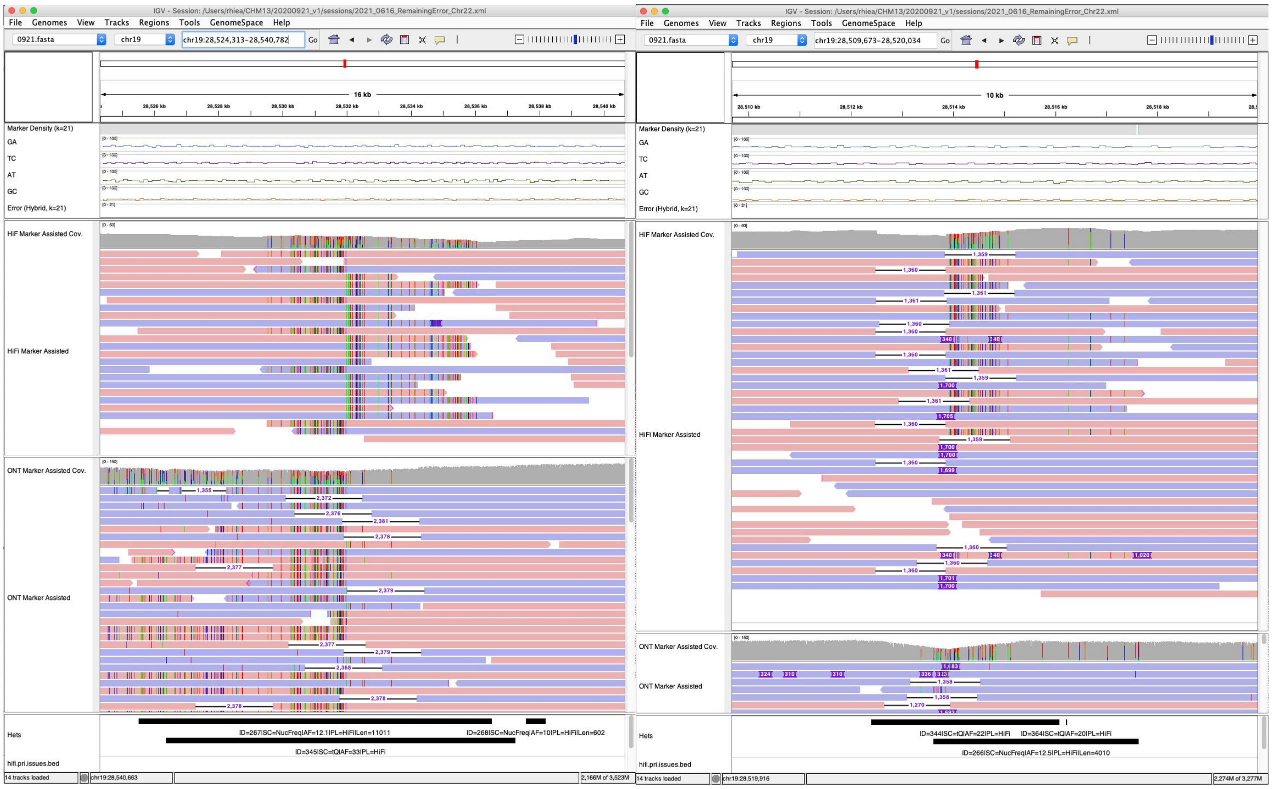


**Supplementary Fig. 11| Chimeric junction of two haplotypes.** In the shown above regions, both HiFi and ONT reads indicate that the consensus has a chimeric junction of the two haplotypes.

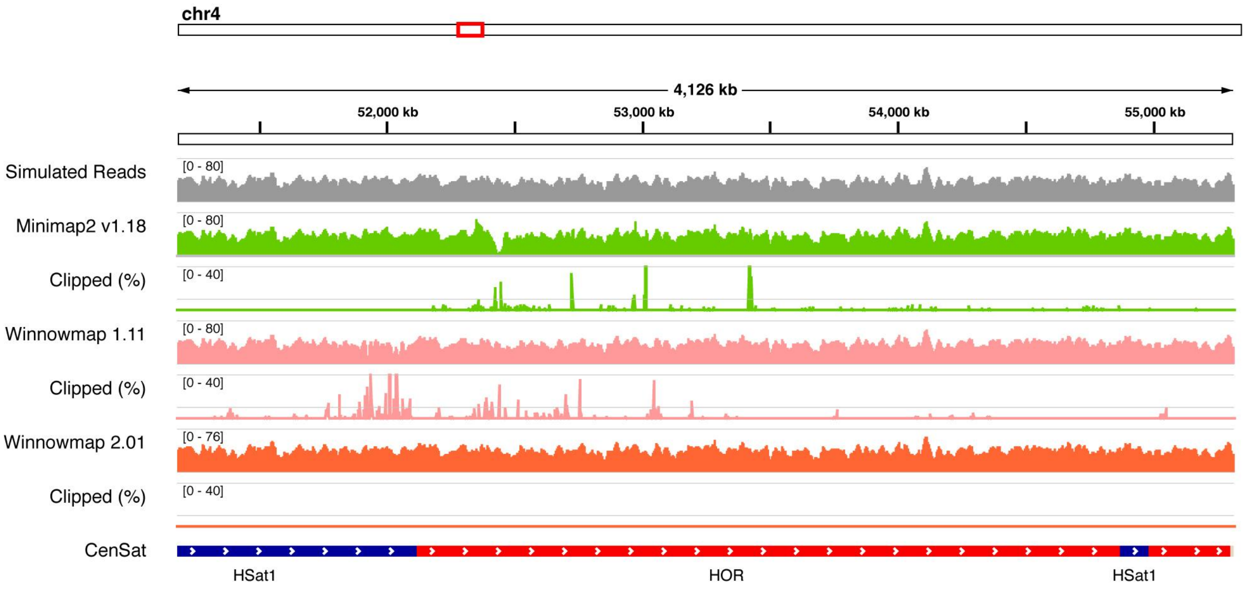


**Supplementary Fig. 12 | Mapping biases found and corrected.** On simulated HiFi reads, we found excessive clippings in highly identical satellite repeats in Minimap and Winnowmap by the time of evaluation. We have addressed this issue in Winnowmap 2.01+. Clipped (%) indicates the percentage of reads clipped in every 1024 bp window, shown in 0~40% range with a midline of 10%.

**Supplementary Table 1. K-mer based consensus quality evaluation.** From each sequencing dataset and assembly versions, 21-mers were collected and compared with Merqury.

| | PacBio HiFi | Illumina | Hybrid |
|---|---|---|---|
| QV | | | |
| V0.9 | 69.68 | 66.09 | 70.22 |
| V1.0 | 69.88 | 67.28 | 72.62 |
| V1.1 | 69.80 | 67.86 | 73.94 |
| K-mers found only in assembly (Error k-mers) | | | |
| V0.9 | 6,881 | 15,723 | 6,073 |
| V1.0 | 6,581 | 11,961 | 3,496 |
| V1.1 | 6,724 | 10,497 | 2,591 |
| K-mers found in both assembly and reads | | | |
| V0.9 | 3,045,438,411 | 3,045,438,411 | 3,045,438,411 |
| V1.0 | 3,045,440,942 | 3,045,440,942 | 3,045,440,942 |

**Supplementary Table 2 | Remaining issues identified from both PacBio HiFi and ONT**

| CHM13 v1.0 | | | CHM13 v1.1 | | | |
|---|---|---|---|---|---|---|
| Chr. | Start | End | Chr. | Start | End | Reason |
| 11 | 54243938 | 54285164 | 11 | 54258631 | 54260218 | Low consensus quality |
| 16 | 36735111 | 36775613 | 16 | 36753902 | 36757498 | Low consensus quality |
| n/a | | | 15 | 3642490 | 3643193 | GA sequencing bias in model rDNA sequence |
| n/a | | | 15 | 3732691 | 3732910 | GA sequencing bias in model rDNA sequence |
| 22 | 39136546 | 39145169 | 22 | 39107778 | 39116401 | Collapsed low complexity sequence |
| 19 | 28509243 | 28519604 | 19 | 28509246 | 28519607 | Chimeric consensus of two haplotypes |
| 19 | 28527178 | 28538917 | 19 | 28527181 | 28538920 | Chimeric consensus of two haplotypes |

**Supplementary Table 3. Low coverage regions detected only from HiFi alignments.** Regions with <7x Winnowmap primary read alignments were collected and categorized given the mapping quality, alignment identity, and sequence context (% microsatellites within 10 kb).

| Cause of low coverage | Num. of regions | (%) |
|---|---|---|
| Likely low consensus qual. | 11 | 5.0% |
| Low consensus qual. | 30 | 13.8% |
| AT biases | 7 | 3.2% |
| GA or TC biases | 170 | 78.0% |
